# HP1β Chromo Shadow Domain facilitates H2A ubiquitination for BRCA1 recruitment at DNA double-strand breaks

**DOI:** 10.1101/2022.01.11.475809

**Authors:** Vijaya Charaka, Sharmista Chakraborty, Chi-Lin Tsai, Xiaoyan Wang, Raj K Pandita, John A. Tainer, Clayton R Hunt, Tej K. Pandita

**Affiliations:** Department of Radiation Oncology, The Houston Methodist Research Institute, Houston TX 77030, USA; Department of Molecular and Cellular Oncology, The University of Texas MD Anderson Cancer Center, Houston, TX 77030, USA; Department of Molecular and Cellular Biology, Baylor College of Medicine, Houston, TX 77030, USA; Center for Genomics and Precision Medicine, Texas A&M College of Medicine, Houston, Texas, USA

**Keywords:** HP1β, chromo shadow domain, BRCA1, ubiquitination, homologous recombination

## Abstract

Efficient DNA double strand break (DSB) repair by homologous recombination (HR), as orchestrated by histone and non-histone proteins, is critical to genome stability, replication, transcription, and cancer avoidance. Here we report that Heterochromatin Protein1 beta (HP1β) acts as a key component of the HR DNA resection step by regulating BRCA1 enrichment at DNA damage sites, a function largely dependent on the HP1β chromo shadow domain (CSD). HP1β itself is enriched at DSBs within gene-rich regions through a CSD interaction with Chromatin Assembly Factor 1 (CAF1) and HP1 β depletion impairs subsequent BRCA1 enrichment. An added interaction of the HP1 β CSD with the Polycomb Repressor Complex 1 ubiquitinase component RING1A facilitates BRCA1 recruitment by increasing H2A lysine 118-119 ubiquitination, a marker for BRCA1 recruitment. Our findings reveal that HP1β interactions, mediated through its CSD with RING1A, promote H2A ubiquitination and facilitate BRCA1 recruitment at DNA damage sites, a critical step in DSB repair by the HR pathway. These collective results unveil how HP1β is recruited to DSBs in gene-rich regions and how HP1β subsequently promotes BRCA1 recruitment to further HR DNA damage repair by stimulating CtIP-dependent resection.

## Introduction

In eukaryotes, genomic DNA is associated with histone and non-histone proteins, that provide the chromatin organization required to regulate transcription, replication, recombination, and DNA repair (Finn and Misteli, 2019; Hunt et al., 2013; Kumar et al., 2012; Pandita and Richardson, 2009; Serizay and Ahringer, 2018). Disturbances in chromatin structure and remodeling are, therefore, often associated with cancer development and progression (Singh et al., 2000; Zhao et al., 2021). Chromatin structure is modulated further by post-translational modifications to the associated proteins (phosphorylation, acetylation, and ubiquitination among others). These modifications can dynamically regulate subsequent recruitment of factors required to initiate a wide range of molecular processes, including the proteins necessary for repair of DNA damage sites (Bennett et al., 2013; Georgoulis et al., 2017; Guo et al., 2018; Hunt et al., 2013; Kaidi et al., 2010; Li et al., 2018; Mendez-Acuna et al., 2010; Shima et al., 2013). At DNA damage sites, histone H2AX is rapidly phosphorylated (γ-H2AX) to act as a marker signaling DNA damage and also aids repair factor recruitment (Horikoshi et al., 2016; Mujoo et al., 2017; Scully and Xie, 2013; Williams et al., 2010; Xie et al., 2010; Yuan and Chen, 2010). Preexisting chromatin modifications also play an important role in the DNA damage response (DDR), for example methylation at H4K20 is essential for 53BP1 foci formation (Kakarougkas et al., 2013; Svobodova Kovarikova et al., 2018) and H4K16ac in transcribing regions is critical for DSB repair by homologous recombination (Horikoshi et al., 2019). Methylation at H3K4 and H3K79 in transcribing regions drives non-homologous end joining (NHEJ) mediated DNA double strand break (DSB) repair (Wei et al., 2018). H3K36 methylation is implicated in DNA repair pathway choice by enhancing the association of the MRN complex to promote NHEJ as well as recruitment of NBS1 through its BRCT2 domain (Fnu et al., 2011).

Methylated histone H3K9 serves as a binding site for isoforms of HP1 (HP1α, HP1β and HP1γ) that are important for heterochromatin formation. All three HP1 isoforms have a related overall structure, an N-terminal chromodomain (CD) sequence, which is responsible for H3K9me2 binding, connected by a hinge region to a variable C-terminal chromoshadow domain (CSD). Depletion of HP1β induces genomic instability (Aucott et al., 2008), though the mechanism is unclear. After DSB formation, HP1β or H3K9me2 are enriched at the break sites and help recruit additional DNA compacting proteins, like suv39H1, which further increase H3K9me2 and HP1β around the sites, leading to gene silencing (Chaturvedi et al., 2012). Moreover, HP1β also plays an important role in DSB repair by HR, as HP1β depletion decreased repair foci formation (Ayoub et al., 2008; Ball and Yokomori, 2009) including those foci containing the important HR repair factor BRCA1 (Lee et al., 2013). Here we show that following DSB induction, HP1β recruits RING1A which ubiquitinates H2A at lysine 118 and 119, a histone modification needed for BRCA1 recruitment and resection of the DSB site prior to repair by HR.

## Results

### Radiation sensitivity induced by HP1β depletion is reversed by expression of the HP1β chromoshadow domain (CSD) alone

To analyze the contribution of individual HP1β domains to a functional DDR, we expressed CSD HP1β (Flag-CSD HP1β, Δ amino acids 22-79) or CD HP1β (Flag-CD HP1β, Δ amino acids to 117-185) regions (**Fig. 1A**) in H1299 cells after depletion of endogenous HP1β by UTR specific siRNA (**Fig. 1B**). Depletion of HP1β increased cell killing by IR, and was rescued by expression of full length Flag HP1β or Flag-CSD but not Flag-CD (**Fig. 1C**). Depletion of HP1β had minimum effect on the initial appearance of IR-induced γ-H2AX and 53BP1 foci; however, residual foci after 4 or 8 hr post-irradiation were higher in HP1β depleted cells than in control cells which was rescued by expression of CSD but not CD (**Fig. 1D,E, Supplementary Fig. 1A,B)**. Higher residual γ-H2AX and 53BP1 foci after DNA damage suggests defective DSB repair (Gupta et al., 2014a; Kumar et al., 2012; Ward et al., 2003). Since radiation damage can be repaired by either NHEJ or HR, we determined which DNA repair pathway(s) were sensitive to HP1β depletion.

**Fig 1.**
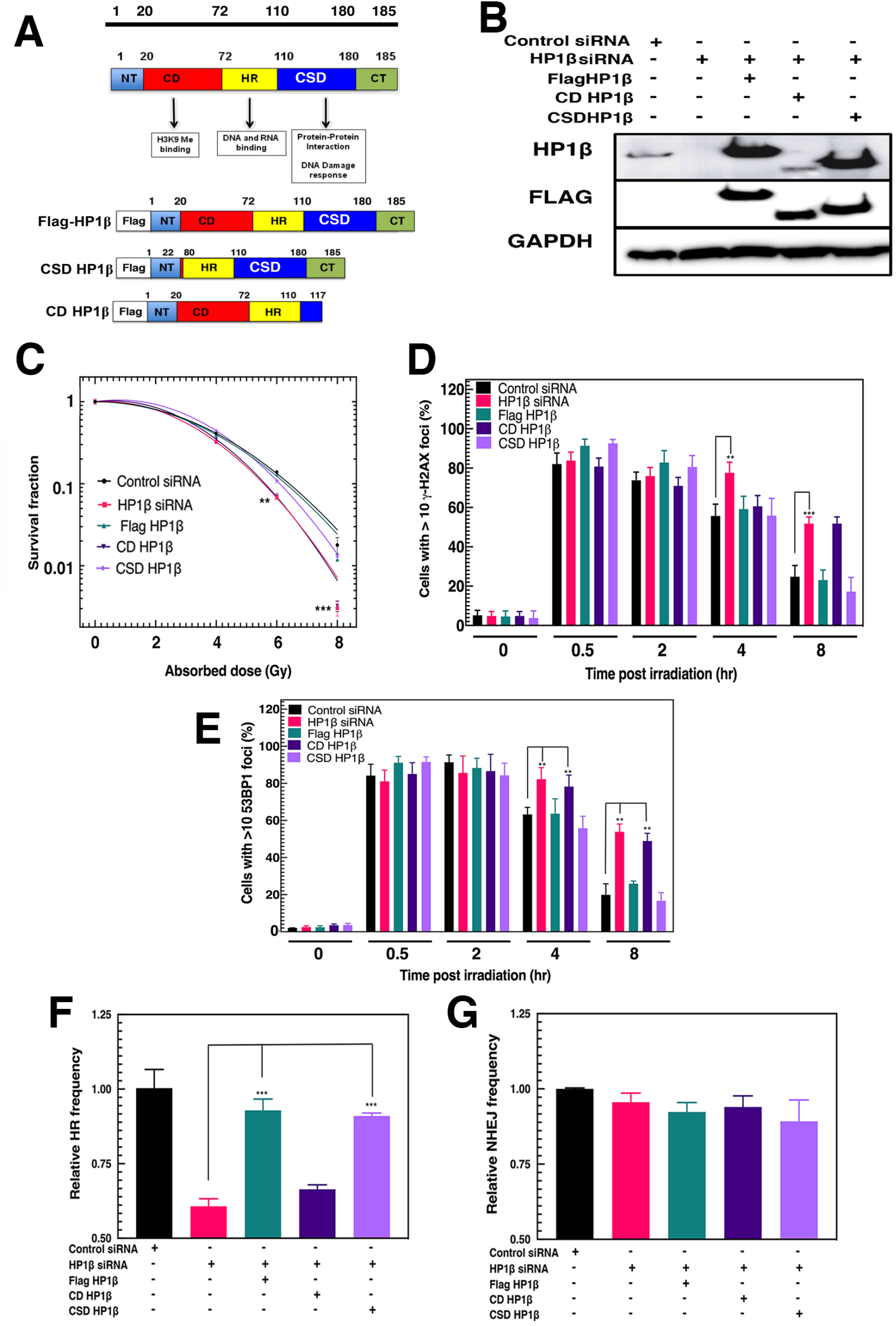
The HP1β chromo shadow domain (CSD) contributes to HR mediated repair of IR-induced DNA damage and cell survival. A) Map of HP1β showing the chromo domain (20 aa-72 aa), hinge region (72 aa – 110 aa) and chromo shadow domain (110aa-180aa). HP1β mutant CD HP1β was generated by deleting amino acids 117 to 185 and CSD HP1β was generated by deleting amino acids 22 to 79 from HP1β. B) Transfected HP1β gene expression in H1299 cells after endogenous HP1β depletion with a 3’ UTR-HP1β siRNA (pCDNA3.1 FLAG HP1β, pCDNA3.1 FLAG CD HP1β, pCDNA3.1 FLAG CSD HP1β). Cellular extracts are prepared 48hr after transfection and FLAG HP1β, CD HP1β, CSD HP1β determined by western blotting. C) Cell survival after increasing IR doses of HP1β depleted cells, ectopically expressing Flag HP1β, CD HP1β, or CSD HP1β as measured by colony formation assay D) Delayed resolution of IR-induced (2Gy) γ-H2AX foci, detected by immuno fluorescent staining, in HP1β depleted H1299 cells is rescued by Flag HP1β or CSD but not CD HP1β expression. E) Dissolution of IR-induced, 53BP1 foci is delayed by HP1β depletion and restored by ectopic Flag HP1β or CSD expression F) Defective HR repair in HP1β depleted DR-GFP cells is restored by ectopic Flag HP1β or CSD expression G) HP1β depletion does not affect DNA DSB repair by NHEJ as measured in Ch1B–EJ5 cells. All standard deviations were calculated from three independent experiments. ^**^*p* < 0.01; ^***^*p* < 0.001.

We ectopically expressed wild-type or HP1β mutants in DR-GFP and EJ5-CHIB expressing cell lines (Horikoshi et al., 2019) previously depleted of HP1β. Depletion of endogenous HP1β reduced DSB repair by HR as measured in DR-GFP cells but expression of either wild type HP1β or the CSD domain alone rescued HR mediated repair. Expression of the CD HP1β domain, however, did not rescue HR mediated DSB repair (**Fig. 1F**). Measurement of NHEJ repair in EJ5-CHIB cells indicated HP1β depletion had no significant effect on repair mediated by this pathway (**Fig. 1G**). The EJ5-CHIB cell assay tests NHEJ at a single, integrated genomic site but identical results were obtained after transient transfection of a linearized NHEJ substrate plasmid template into endogenous HP1β depleted cells (**Supplementary Fig. 1C**), further confirming HP1β depletion does not impair DSB repair by NHEJ.

### CSD HP1β recruitment at DNA damage sites involves CAF1

During the early DDR response after DNA strand breakage, HP1β protein is recruited to break sites to promote H3K9 methylation (Ayrapetov et al., 2014; Zeng et al., 2010). Several reports have shown that HP1β binding to H3K9me2 increases retention of KAP1, and HP1β along with suv39h1 ensure gene silencing during DSB repair (Ayrapetov et al., 2014; Sharma et al., 2003). To examine HP1β recruitment near DNA break sites, three I-Sce1 inducible DSB sites were inserted at different specific positions in the genome (Chr1A, Chr1B and Chr1C) as described (**Supplementary Fig. 2A**) (Horikoshi et al., 2019) and protein levels at the DSB sites analyzed by ChIP/qPCR (Horikoshi et al., 2019). The Chr1A and Chr1C I-Sce1 sites are in gene rich and H4K16ac high density regions while the Chr1B site is in a gene and H4K16ac poor region (Horikoshi et al., 2019). As expected, both γ-H2AX and HP1β increased at the Chr1A, Chr1B and Chr1C sites after DSB induction (**Fig. 2A, B**), but a relatively higher fold enrichment of HP1β in H4K16ac rich regions was observed as compared to the H4K16ac poor region (**Fig. 2A**). Analysis of the individual HP1β CSD and CD domains indicated both were highly enriched at DSBs in gene-rich regions, where HR is known to be predominant (Horikoshi et al., 2019) (**Fig. 2C, D**). In contrast, both HP1β mutants displayed only a small increase in levels at the gene poor region Chr1B site after DSB induction (**Fig. 2C, D**).

**Fig 2.**
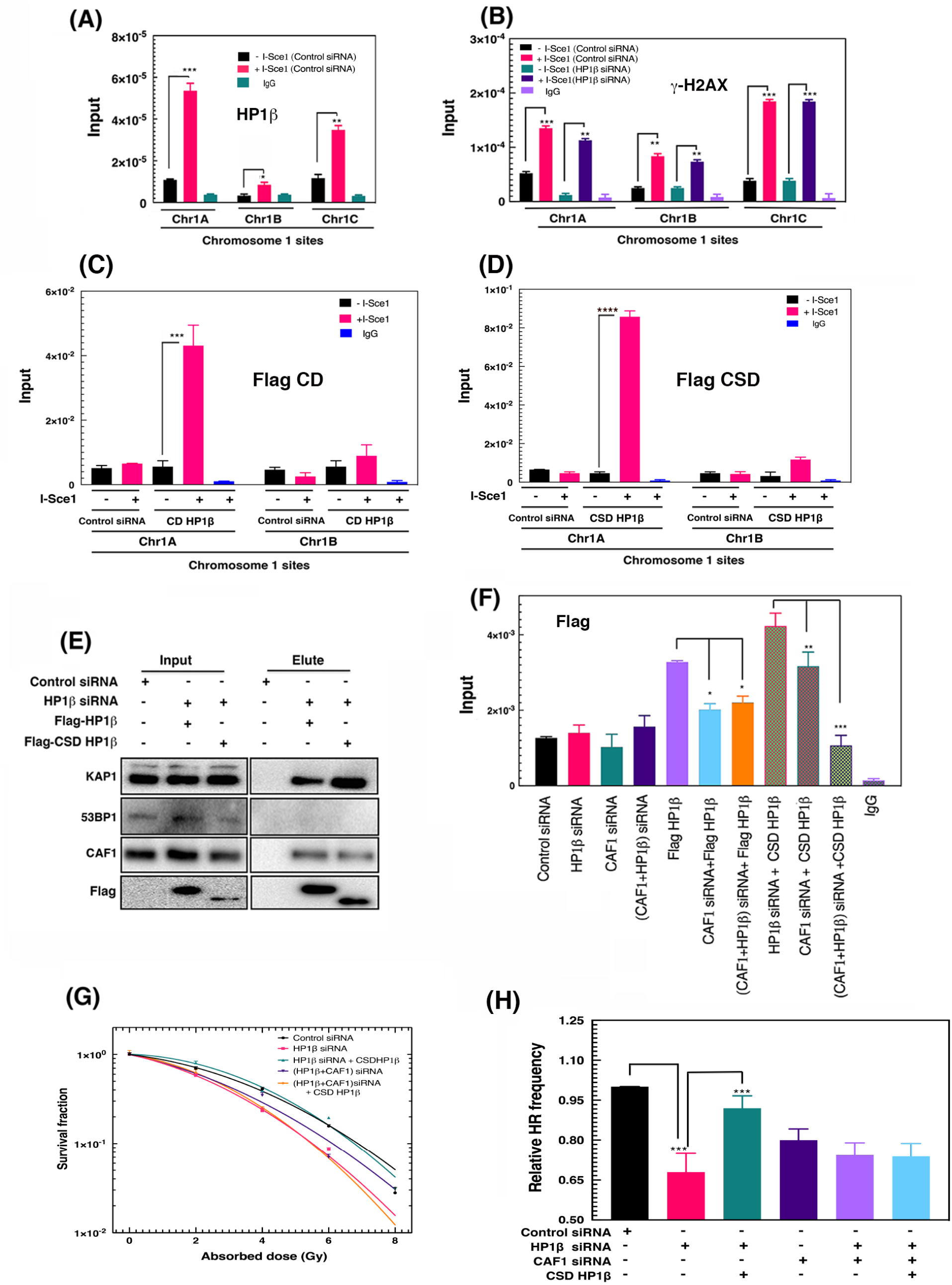
CSD HP1β is recruited to DSB sites through CAF1. A) Increased HP1β recruitment to break sites as measured by ChIP/qPCR at different I-Sce1 induced cleavage sites. B) Depletion of HP1β does not alter DSB induction as measured by γ-H2AX ChIP/qPCR at different ISce1 sites. C&D) Flag CD HP1β and Flag CSD HP1β recruited to Chr1A and Chr1B DSB sites. Using ChIP-q PCR, CD HP1β and CSD HP1β recruitment was determined. E) CAF1 and KAP1 co-immunoprecipitated with Flag HP1β and CSD HP1β. F) CAF1 deletion decreases CSD HP1β and Flag HP1β recruitment at the Chr1A DSB site. G) CAF1 depletion blocks CSD mediated rescue of irradiated HP1β depleted cells as determined by clonogenic cell survival assay. H) Depletion of CAF1 blocks CSD mediated rescue of homologous recombination in HP1β depleted cells as determined by DR-GFP assay. All standard deviations were calculated form at least three independent experiments. \**p* < 0.05; ^**^p < 0.01; ^***^*p* < 0.001.

Previous reports found that repressive marker (HP1β and H3K9me2) enrichment at DNA break sites promoted additional recruitment/marks around the site (Alagoz et al., 2015). We found that H3K9me2 levels increased after DNA damage, but HP1β loss reduced the H3K9me2 levels at the break site (**Supplementary Fig. 2B**).

Luijsterburg and co-workers suggested that HP1β recruitment is independent of both its CD, which is the methyl-binding domain, and chromatin H3K9me2 (Luijsterburg et al., 2009). We also found that while CSD HP1β became enriched, the HP1β methyl-binding CD domain was not enriched at DSB sites in transcribed regions, another indication HP1β recruitment is H3K9 methylation independent (**Fig. 2C, D**). To determine whether any known CSD interacting proteins impacted HP1β recruitment, we performed co-immunoprecipitation studies and confirmed that both KAP1 and CAF1 interact with HP1β (Bartova et al., 2017; Brasher et al., 2000; Lemaitre and Soutoglou, 2014; Lorkovic et al., 2017; Nakatani et al., 2006; White et al., 2012), more specifically the CSD (**Fig. 2E, Supplementary Fig. 2C**).

CAF1 is involved in histone assembly, we next determined whether CAF1 depletion altered HP1β recruitment at DNA damage sites. ChIP-based recruitment analysis indicated that CAF1 depletion decreased both HP1β and CSD HP1β recruitment to DSBs (**Fig. 2F**). Moreover, CSD HP1β expression after CAF1/HP1β co-depletion no longer rescued the IR sensitive phenotype of HP1β depleted cells (**Fig. 2G**). In addition, DNA DSB repair by HR was no longer rescued in HP1β depleted cells by CSD HP1β expression in CAF1 co-depleted cells (**Fig. 2H**). In contrast, CSD HP1β did restore HR in KAP1/HP1β co-depleted cells (**Supplementary Fig. 2D**), suggesting HP1β CSD interaction with CAF1, but not KAP1, is critical for effective HR repair.

### The HP1β CSD and DNA end resection

Depletion of HP1β has been reported to decrease BRCA1 foci formation (Alagoz et al., 2015; Fukuda et al., 2016; Lee et al., 2013), which is essential for DNA resection; however, details of the underlying mechanism are unclear. During DNA damage repair, displacement of 53BP1 at DSBs requires BRCA1 and occurs prior to recruitment of proteins involved in resection (Feng et al., 2013; Mattoo et al., 2017). To identify a possible role for HP1β in resection, we measured single strand DNA production at a DSB site in control and HP1β depleted cells using the ER-AsiSl assay system (Zhou et al., 2014) (**Fig. 3A, Supplementary Fig. 3A**). Time course studies in control cells of the tamoxifen induced ER-AsiSI DNA cleavage site indicated single stranded DNA (ssDNA) levels started to peak from 2hr and maximum ssDNA formation occurred at 4hr with a slight decline by 6hr (**Fig. 3A**). As expected, measured resection at the distal site 1618 bp from the DSB was proportionately slightly less than at the proximal 335 bp site. In contrast, after HP1β depletion, ssDNA formation was substantially less and only slowly increased before reaching a maxima at 6 hr with significantly decreased formation 335 bp or 1618 bp from the DSB site (**Fig. 3A**).

**Fig 3.**
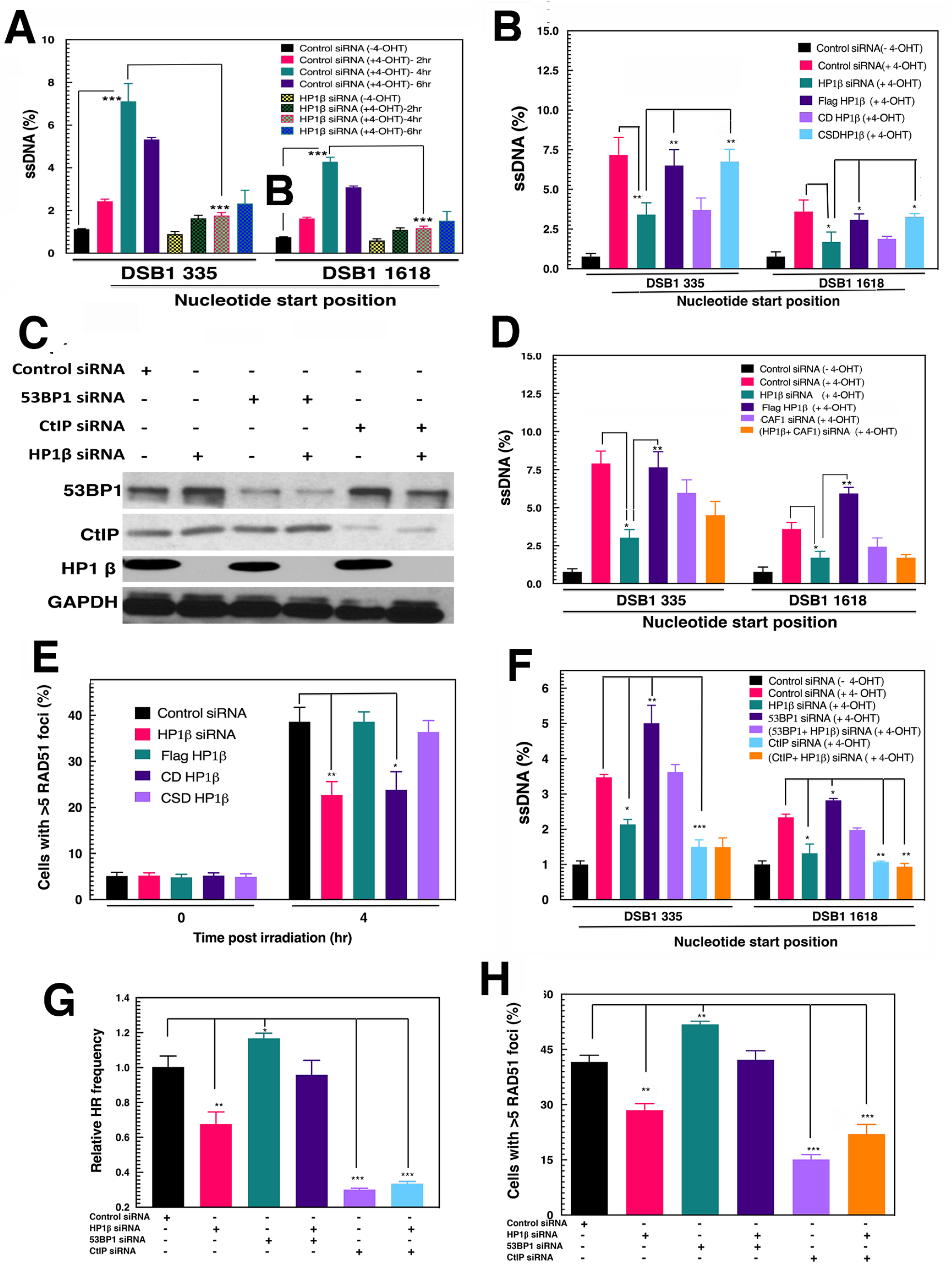
DNA resection at DSBs, Rad51 foci formation and HR repair in HP1β depleted cells are rescued by CSD HP1β expression. A) Decreased ssDNA generated at DSBs in HP1β depleted cells as measured at the 335 bp and 1618 bp after DSB induction with tamoxifen. Amount of after resection was measured at 335 and 1618 bp 3’ from the DSB site using two different primer sets for qPCR. SD values were determined in between three independent experiments. ^*^*p* < 0.05; ^**^*p* < 0.01; ^***^*p* < 0.001. B) Single strand DNA formation at a DSB in HP1β depleted cells ectopically expressing CD HP1β or CSD HP1β was measured proximal and distal to the site by qPCR using the ER-AsiSI system as described in Material and Methods. C) Western blot showing knockdown of CAF1 and CtIP by specific siRNA. D)Percentage of single strand DNA formed in CSD-HP1β expressing cells was quantitated in presence and absence of CAF1. E) RAD51 foci formation in irradiated HP1β depleted cells is restored by CSD but not CD expression. Approximately 300 cells with RAD51 foci were examined and standard deviation from three experiments was calculated. F) 53BP1 depletion does not rescue HP1β dependent DNA resection. ssDNA generated after resection was measured at proximal and distal sites to a DSB using qPCR in cells with and without. G) DR-GFP cells were transfected with respective siRNA’s (Control siRNA, HP1β siRNA, 53BP1 siRNA, and CtIP siRNA) and after 48hr cells were re-transfected with I-Sce1 plasmid. The percentage of GFP positive cells was measured by using FACS. Percentage of GFP positive cells was normalized to control cells. H) Cells with and without above-mentioned siRNA were enumerated for RAD51 foci after irradiation. The three independent experiments were performed and standard deviation between these experiments was calculated. ^*^*p* < 0.05; ^**^*p* < 0.01; ^***^*p* < 0.001

Expression of CSD HP1β, but not CD HP1β, restored ssDNA formation to levels in non-HP1β depleted cells (**Fig. 3B**). Since CAF1 interacts with HP1β (**Fig. 2E)**, we further examined whether depletion of CAF1 also affected resection and found CAF1 depletion blocked the ability of CSD expression to restore resection in HP1β depleted cells (**Fig. 3C, D**). Consistent with these results, foci formation by the DNA single strand binding protein RAD51 was reduced in HP1β depleted cells and restored by HP1β or CSD HP1β expression (**Fig. 3E, Supplementary Fig. 3B**). These results indicate that HP1β is required at an early step in HR-mediated DNA repair (**Fig. 3A-C**) that is functionally dependent on the HP1β CSD.

Furthermore, we observed that 53BP1 depletion increased single strand DNA (ssDNA) and RAD51 foci formation but concurrent HP1β depletion restored normal levels of ssDNA and RAD51 foci formation (**Fig. 3E-G**). CtIP depletion, as expected, decreased ssDNA and RAD51 foci formation, and this was not further decreased by simultaneous HP1β depletion (**Fig. 3E, F**), suggesting that CtIP and HP1β function in the same pathway for resection. These results correlated with DSB repair by HR (**Fig. 3F, G, Supplementary Fig. 3C**) in that HP1β or CtIP depletion reduced HR-mediated DSB repair.

### CSD HP1β is critical for 53BP1 displacement and BRCA1 foci formation

HP1β depletion reduced the frequency of IR-induced BRCA1 foci formation. Since 53BP1 and RIF1 block IR-induced BRCA1 foci formation (Bunting et al., 2010; Escribano-Diaz et al., 2013), we expressed wild type or HP1β mutants in HP1β depleted cells and measured co-localization of 53BP1 and RIF1 foci after irradiation (**Supplementary Fig. 3D**). Neither HP1β depletion nor expression of CD HP1β or CSD HP1β in depleted cells affected IR-induced 53BP1/RIF1 co-localization (**Supplementary Fig. 3D**). However, HP1β depletion reduced the frequency of BRCA1 foci formation (**Fig. 4A, B**). To determine which domain of HP1β is required for BRCA1 recruitment at DNA damage, we depleted endogenous HP1β and expressed wild type, CD or CSD domain. We found that wild-type and CSD HP1β expression, but not CD HP1β,rescued BRCA1 foci formation (**Fig. 4A, B**). HP1β depleted cells had reduced co-localization of 53BP1/BRCA1 foci, whereas CSD HP1β expression rescued co-localization of 53BP1/BRCA1, but not CD expression (**Fig. 4A-C**).

**Fig 4.**
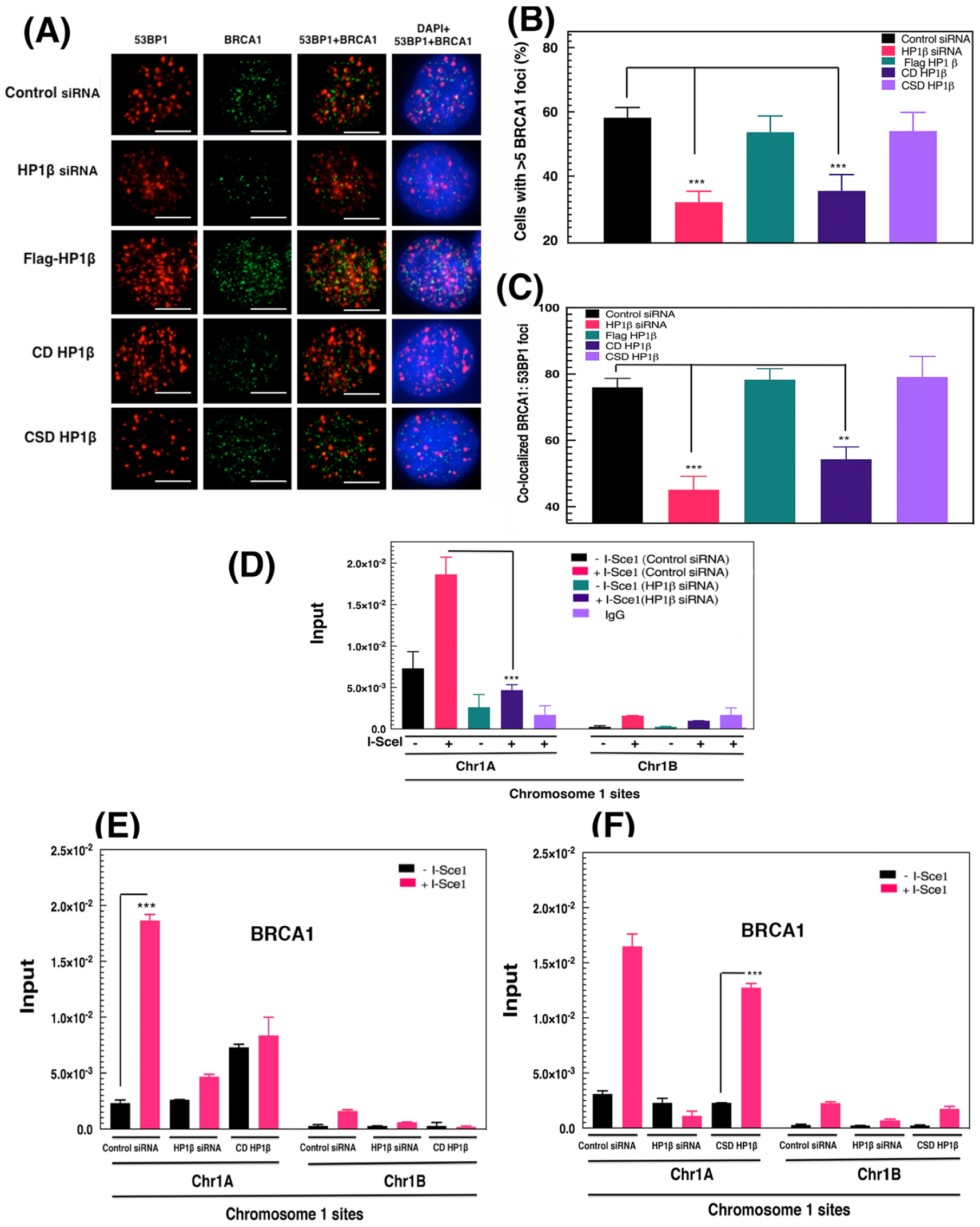
CSD HP1β promotes BRCA1 recruitment at DSB sites. A) Immuno-fluorescent detection of BRCA1 and 53BP1 foci in irradiated, HP1β depleted H1299 cells expressing FLAG HP1β or FLAG domain mutants. Images were captured on an Axio-Vision Imager 2 microscope for analysis B) BRCA1 foci measured 6h after irradiation in cells treated as in A. C) Co-immunostaining for 53BP1 and BRCA in post-irradiated fixed samples. Co-localized 53BP1: BRCA1 foci were counted in 3 sets of 50 cells, and the percent co-localized 53BP1/BRCA1 foci was calculated relative to total number of foci (53BP1 plus BRCA1). D) HP1β depletion decreases BRCA1 recruitment to DSB sites. BRCA1 levels at the Chr1A and Chr1B sites were measured by ChIP-qPCR before and 20h after I-Sce1 transfection. E&F) BRCA1 recruitment at the Chr1A and Chr1B DSB sites was measured in HP1β depleted cells with or without FLAG CD HP1β (E) or FLAG CSD HP1β (F) expression. Standard deviations calculated from a minimum of three independent experiments. ^**^*p* < 0.01; ^***^*p* < 0.001, Scale bar is 10 μm.

To directly measure BRCA1 recruitment at DSBs, we used two different I–Sce1 inducible DSB sites cell lines (Chr1A, Chr1B) as described previously (Supplementary Fig. 2) (Horikoshi et al., 2019). Chromatin immuno-precipitation (ChIP) studies detected a significant increase in BRCA1 at the Chr1A site (gene-rich) after DSB induction and an insignificant BRCA1 increase at the Chr1B site (gene-poor) (**Fig. 4D**). Depletion of HP1β significantly diminished BRCA1 recruitment at the Chr1A DSB site (**Fig. 4D-F**). The ability of the HP1β deletion mutants to restore BRCA1 recruitment to DSBs differed dramatically, as CD HP1β expression did not restore BRCA1 recruitment in HP1β depleted cells (**Fig. 4E**), whereas CSD HP1β restored BRCA1 recruitment (**Fig. 4F**).

### The influence of HP1β on cell survival after irradiation is ATM independent

To determine whether the HP1β repair function is ATM dependent, HP1β depleted cells were treated with an ATM inhibitor (KU55993) and cell survival measured after IR (**Fig. 5A**). Either HP1β depletion or ATM inhibition increased cell sensitivity to IR (**Fig. 1B, 5A**). The combined treatments produced a further, synergistic decrease in cell survival (**Fig. 5A**), suggesting that HP1β and ATM effect survival by separate pathways. This was further confirmed by depletion of HP1β in ATM null cells (GM5823+hTERT) (Wood et al., 2001) where depletion of HP1β (**Supplementary Fig. 4A**) further decreased cell survival post irradiation (**Supplementary Fig. 4B**). Depletion of HP1β had no effect on IR-induced ATM auto-phosphorylation (**Supplementary Fig. 4B,C**) and phosphorylated p-MDC1 foci formation (**Fig. 5B, Supplementary Fig. 4D**), suggesting HP1β functions independently of the ATM-mediated pathway in the IR-induced DNA damage response.

**Fig 5.**
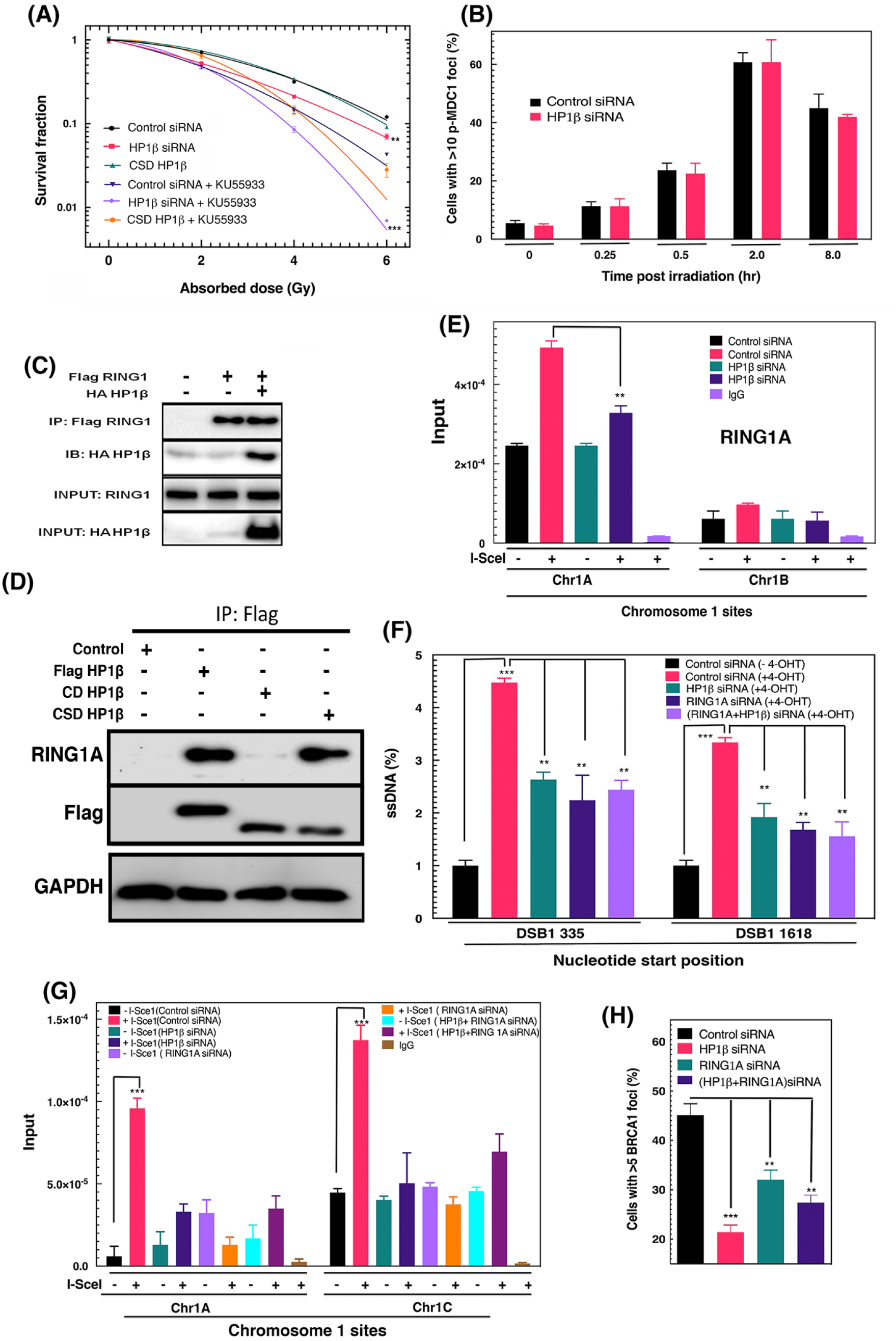
HP1β interacts with RING1A and enhances BRCA1 recruitment at DSB sites. A) Effect of ATM inhibition on clonogenic survival of irradiated cells. Cells were treated with 10μM ATM inhibitor (KU-55933) for 1hr before IR treatment. B) p-MDC1 foci formation/dissolution in irradiated H1299 cells is not altered by HP1β depletion. C) FLAG RING1A co-immunoprecipitated with HA HP1β. D) Flag RING1A interaction with CD HP1β and CSD HP1β E) RING1A enrichment after DNA damage at Chr1A and Chr1B DSB sites as measured by ChIP-qPCR is decreased by HP1β depletion. F) Decreased single strand DNA formation at DSBs in RING1A or HP1β depleted cells as measured by the ER-AsiSI assay G) Impact of RING1A depletion on BRCA1 recruitment at Chr1A and Chr1C DSB sites before and after I-Sce1 transfection H) Effect of RING1A depletion on HR frequency in DR-GFP cells with and without HP1β. Experiments were done for three times and Standard deviation was calculated. ^**^*p* < 0.01; ^***^*p* < 0.001

### HP1β interacts with RING1A to facilitate BRCA1 foci formation

Mono-ubiquitinated H2A at lysine 119 is essential for BRCA1 recruitment to DSBs (Hung et al., 2017). We therefore tested for interactions between HP1β and Polycomb repressive complex 1 (PRC1), which contains an E3 ubiquitin ligase, by co-immunoprecipitation. We found that HP1β interacts with the PRC1 complex protein RING1A (**Fig. 5C,D**). Deletion analysis indicated HP1β interacted with RING1A specifically through CSD but not CD (**Fig. 5D**).

HP1β/RING1A interaction increases substantially in cells 30 mins after irradiation (**Supplementary Fig. 5A**). RING1A was enriched after DSB induction at the gene-rich region (Chr1A), while less enrichment was observed at the gene poor DSB site (**Fig. 5E**). This interaction was confirmed by in vitro studies using recombinantly purified RING1A-FL and CSD HP1β where CSD HP1β pulls down RING1A-FL (**Supplementary Fig. 5B**).

Additional insights into the HP1β and RING1A interaction were obtained from protein docking models. Full-length HP1β and RING1A were modelled using TrRosetta (Yang et al., 2020) before running the rigid-body protein-protein docking using ClusPro docking server (Kozakov et al., 2017). The selection criteria of top-ranked model were based on the known protein-protein interactions on RING1A, where its N-terminal domain interacts with polycomb complex protein Bmi1 (McGinty et al., 2014), and on HP1β, where the CSD tends to form a dimer via its α -helix (Ekblad et al., 2005; Kumar and Kono, 2020). For example, the I161E mutant on the CSD α-helix was reported to disrupt protein-protein interaction (Brasher et al., 2000). The docking model revealed that the same α-helix of CSD may interact with RING1A (**Supplementary Fig. 5C**). Although CSD HP1β and CD HP1β share a similar fold, their sequence identity is only ∼23%. Sequence conservation and secondary structure analyses show the CSD α-helix is mostly conserved, while CD α-helix is not (**Supplementary Fig. 5D, E**). Moreover, CD HP1β could not bind to RING1A via the same CSD binding site as its predicted long α-helix would seriously clash with the RING1A structure (**Supplementary Fig. 5E**). Interestingly, the RING1A α-helix binding site is also highly conserved, suggesting important functional roles of these residues. In sum, our structure-based docking model predicts the potential RING1A binding site can only interact with the HP1β CSD, not the CD, consistent with our cellular results (**Fig.5D, Supplementary Fig.5**).

To determine whether RING1A promotes BRCA1 recruitment to DSBs, we examined BRCA1 enrichment at the Chr1A and Chr1C sites after DSB induction in cells depleted of either HP1β or RING1A or in combination. We found under all conditions significantly reduced BRCA1 recruitment (**Fig. 5G**). Similarly, RING1A depletion reduced IR-induced BRCA1 foci formation (**Fig. 5H**), and reduced single ssDNA formation 335bp and 1618 bp distal from the DSB (**Fig. 3A, 5F**). These foci and ssDNA reductions resemble those from HP1β depletion, while depletion of both produced no additional decrease (**Fig. 5F**). This indicates HP1β and RING1A function epistatically for resection during DSB repair by HR.

### HP1β promotes ubiquitination of H2A by RING1A

HP1β is required for BRCA1 distribution at DSBs (**Fig. 4**), and the PRC1 complex protein RING1A ubiquitinates H2A at lysine 119 (Benitz et al., 2016; Patel et al., 2017; Rossi et al., 2016). We reasoned, that HP1β might be essential for H2A ubiquitination and subsequent BRCA1 recruitment to DSBs. A significant IR-induced increase in cellular H2A poly-ubiquitination was observed that was largely lost in HP1β depleted cells (**Fig. 6A, B**). Furthermore ubiquitinated (ub)-H2AK119 enrichment was observed at the Chr1A site after DSB induction, whereas HP1β depletion significantly reduced ub-H2AK119 enrichment at the site (**Fig. 6C**).

**Fig 6.**
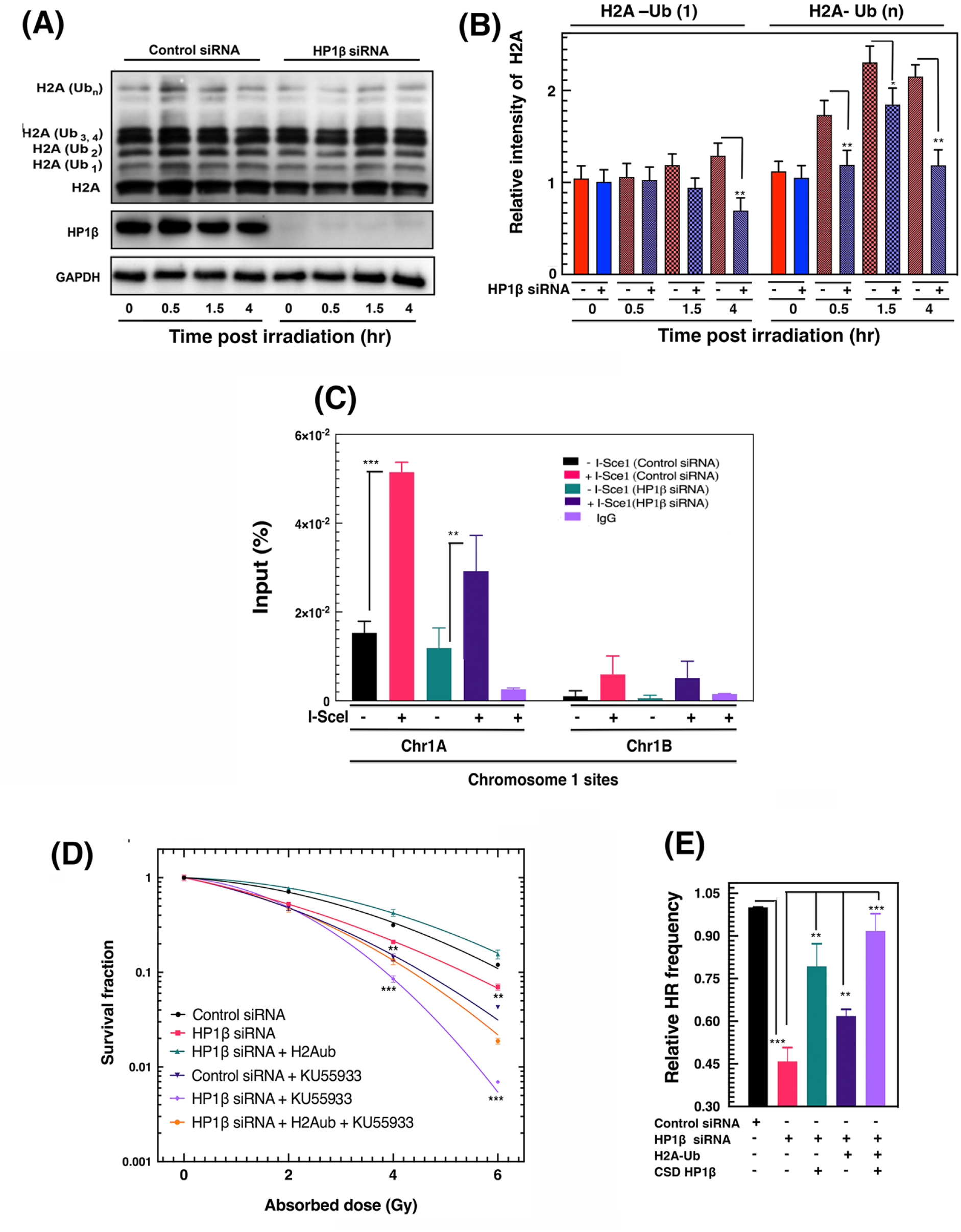
HP1β modulates H2A K119-ub levels at DSB sites. A&B) HP1β depletion decreases H2A poly-ubiquitination. Control and HP1β depleted H1299 cells were transfected with FLAG-H2A vector then irradiated (10 Gy). FLAG immunoprecipitated H2A was analyzed for Ub by western blot analysis with Flag antibody. Relative H2A mono, polyubiquitination at individual time points (0, 0.5, 1.5, 4 hrs) was calculated relative of total H2A. C) Effect of HP1β depletion on H2AK119–ub levels at the Chr1A and Chr1B sites before and after DSB induction as determined by ChIP–qPCR. D) H2A–ub reversal of radio sensitivity in HP1β depleted cells is independent of ATM. Cells with and without HP1β depletion were treated with ATM inhibitor (KU 55933) then treated with increasing IR doses. Effect of H2A-ub expression on cell survival was determined by clonogenic survival. E) Exogenous H2A-ub partially restores HR in HP1β depleted cells as determined by the DR-GFP repair assay. Three independent experiments were conducted and an SD value in between experiments was determined. ^*^*p* < 0.05; ^**^*p* < 0.01; ^***^*p* < 0.001.

These results were complemented by examining the effect of expressing a carboxy-end H2A ubiquitinated protein (Zhu et al., 2011) in HP1β depleted cells.

Expression of the conjugated uH2A protein rescued IR-induced cell killing and improved HR mediated repair in HP1β depleted cells (**Fig. 6D, E**). Expression of ub-H2AK119 rescued cells from enhanced IR-induced cell death whether in the presence or absence of ATM inhibitor, suggesting ub-H2AK119, as does HP1β, functions at DSBs independently of ATM function (**Fig. 6D**). Overall, these results identify an interaction between HP1β and PRC1 complex protein (RING1A) that facilitates H2A ubiquitination and thus BRCA1 recruitment at DSBs (**Fig. 6E**).

## Discussion

Conflicting studies have suggested that HP1β facilitates DNA repair, or is inhibitory to DNA repair (Ayoub et al., 2008; Ayoub et al., 2009; Dinant and Luijsterburg, 2009; Horikoshi et al., 2019). Loss of function HP1β studies in *Caenorhabditis elegans*, yeast and human indicated increased sensitivity to IR, suggesting that HP1β contributes to DNA damage repair by HR (Ahringer and Gasser, 2018; Jang et al., 2018; Wei et al., 2018). Chromatin associated remodeling factors are different in gene rich vs. gene poor regions and can influence the recruitment of different DNA repair factors or proteins at DSB sites (Horikoshi et al., 2019). Interestingly HP1β is more efficiently recruited and promotes HR at DSBs located in gene-rich regions as compared gene-poor regions. Some studies suggest that after DNA damage, HP1β is recruited to DSB sites and then enrichment is extended beyond the break site (Ayrapetov et al., 2014; Goodarzi and Jeggo, 2013; Hunt et al., 2013). However, there are also contradictory results regarding accumulation of HP1β at DSB sites (Ayoub et al., 2009; Ball and Yokomori, 2009; Luijsterburg et al., 2009; Zeng et al., 2010). Adding to the existing literature and to defining the mechanism for HP1β function, we report that HP1β recruitment to DSBs in transcriptionally active regions can be mediated independently by either the HP1β CD or CSD domain; however, CSD HP1β recruitment levels at DSB sites are significantly higher. Further, we explored factors essential for CSD HP1β recruitment at DSB sites and found that CAF1 is required and depletion of CAF1 increased sensitivity to IR (Huang et al., 2018). Reducing CAF1 in HP1β depleted cell blocked the ability of CSD HP1β to rescue the IR sensitivity, indicating CAF1 and CSD HP1β act in the same HR pathway. Consistent with the HP1β role in HR, HP1β depletion reduced BRCA1 foci formation in irradiated cells (Lee and Ann, 2015; Lee et al., 2013; Lee et al., 2015). Earlier studies have shown that BRCA1 plays a role in 5’ - 3’ DNA end resection which generates the ssDNA substrate that is subsequently loaded with RAD51. Importantly, regulation of resection can control DSB repair mechanism, pathway selection, and biological outcomes to DSBs (Dutta et al., 2017; Shibata et al., 2014). Here we report on the mechanism by which BRCA1 recruitment to chromatin DSBs is influenced by HP1β. We found that CSD HP1 β recruits BRCA1 to DSB sites at gene-rich sites. Furthermore, this is independent of ATM as the depletion of HP1β and ATM inhibition increased cell kill synergistically which supports and extends previous results (Alagoz et al., 2015), suggesting that BRCA1 recruitment through CSD HP1β is independent of the ATM function.

Histone ubiquitination modifications (K13-15, K118-119, K127-129) and E3 ubiquitin ligases (RNF8, RING1B, RNF168, BARD1) play critical roles in recruitment of 53BP1 and BRCA1 (Chapman et al., 2012; Daley and Sung, 2014; Munoz et al., 2014). PRC1 complex proteins interact with DNA repair proteins and this affinity is increased after DNA damage (Chandler et al., 2014). In mammalian systems, the PRC1 complexes are heterogeneous with six different PRC1 complexes known (PRC 1.1 to PRC 1.6); however, all these complexes purify together with RING1A/B (Gao et al., 2012). In transcriptionally repressed regions, PRC1 complex proteins ubiquitinate H2A at lysine 118/119, which is required for any subsequent BRCA1 recruitment (Ginjala et al., 2011; Zhu et al., 2011). Direct binding of CSD HP1β with RING1A evidently creates an efficient H2A ubiquitination complex at DSBs in actively transcribing regions since we found that depletion of either HP1β or RING1A reduced H2A K118-119 ubiquitination levels. Moreover, HP1β and RING1A likely act as a complex in BRCA1 recruitment as our results showed no significant difference in H2A-ub118 -119 reduction between depleting HP1β and RING1A separately or together. These collective results unveil how HP1β is recruited to DSBs in gene-rich regions and how HP1β subsequently promotes BRCA1 recruitment to further DNA damage repair during HR by stimulating resection (**Fig. 7**).

**Fig 7.**
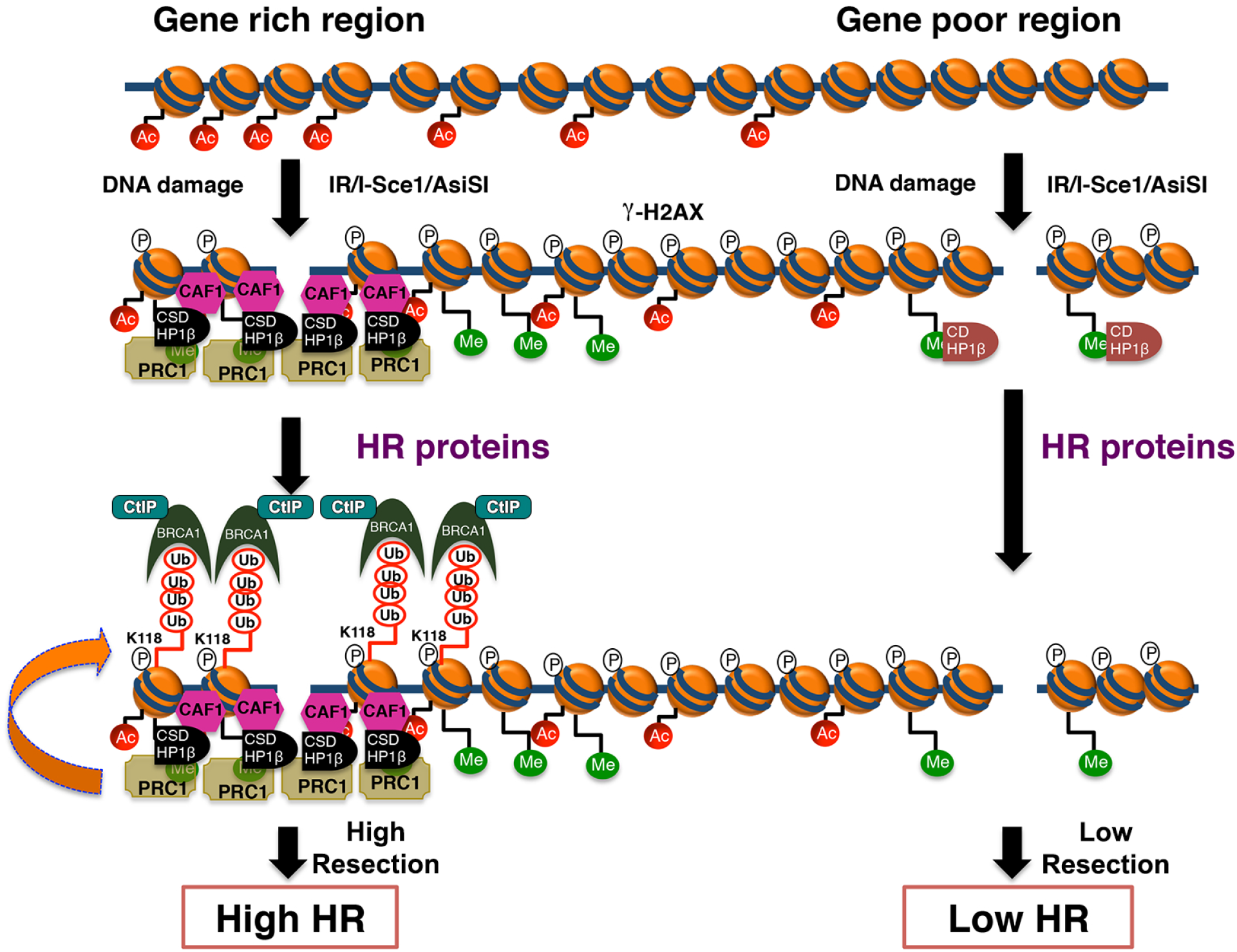
Representative model of HP1β role in BRCA1 recruitment.

## Materials and methods

Human cell lines H1299 and U2OS were obtained from ATCC (Manassas, VA) and cultured in DMEM + 10 % FBS media. Cell transfection with siRNA or plasmid was as described previously(Ahmed et al., 2018; Chakraborty et al., 2018; Mujoo et al., 2017). Resection assay reagents (ER-AsiSI plasmids and U2OS cells) were a generous gift from T. Paull (University of Texas Austin, TX). The pCDNA3.1 FlagHP1β, and mutant DNAs (PCDNA 3.1 CD HP1β and CSD HP1β were transfected into H1299 cells or U2OS cells.

### Antibodies

The list of primary antibodies used for immuno blot (IB or immunofluorescence (IF) analysis are Rat anti-HP1β (IB 1:500,ab10811), mouse anti-HP1γ (IB, 1:1,000;ab217999); mouse anti-HP1α (IB, 1:1,000, ab77256); mouse anti-γH2AX (IB 1:1000, IF: 500, 05-636; Millipore, Mouse anti-human RAD51 (IF, 1:500, ab133534); Rabbit anti RING1A (IB: 1:250, ab32644), Rabbit anti RING2 (IB 1: 500, ab101273) Rabbit anti CtIP (IB: 1:1000, ab70163), Rabbit anti CAF1 (IB: 1000, ab126625), Rabbit anti KAP1 (IB1: 500, ab10484), Rabbit anti 53BP1(IB: 1:500, IF: 1:250, SC-22760) Mouse anti FLAG (IB 1:500), Rabbit anti HA (IB 1:250). Secondary antibodies used were: for IB, HRP-conjugated affinity-purified sheep anti-mouse or donkey anti-rabbit (1:5000; Jackson Immuno Research Laboratories, Inc.); and for IF, goat anti-mouse or anti-rabbit coupled to Alexa Fluor 488 or 594 (1:1,000; Invitrogen).

### Immunofluorescence

H1299 cells were transfected with control siRNA or 3’ UTR HP1β siRNA to deplete endogenous HP1β. HP1β siRNA treated cells were transfected with Flag HP1, Flag CD HP1β, or Flag CSD HP1β. The cells were irradiated and samples obtained at increasing times after irradiation. After fixing with 4% paraformaldehyde or methanol, cells were immuno-stained with different antibodies as mentioned in text (Chakraborty et al., 2018; Gupta et al., 2014b; Horikoshi et al., 2016; Mattoo et al., 2017; Singh et al., 2018). Images captured by using a Zeiss Axio Imager and analyzed by using Image J software.

### ssDNA Quantification assay by using the ER-AsiSI system

A quantitative ssDNA resection product was measured as previously described (Chakraborty et al., 2018; Singh et al., 2018; Zhou et al., 2014). Exponentially growing ER-AsiSI U2OS cells treated with tamoxifen (300 ng/ml) for 3hr, and 6 million treated cells were embedded in 0.6 % low melting agarose. A 50 ul agarose ball of cells was serially treated with 1ml of EDTA Sarcosine proteinase K (ESP) buffer (0.5 M EDTA, 2% N-lauroylsarcosine, 1 mg/ml of proteinase K, and 1 mM CaCl2 [pH 8.0]) and HS buffer ((1.85 M NaCl, 0.15 M KCl, 5 mM MgCl2, 2 mM 213 EDTA, 4 mM MTris, and 0.5% Triton X-100 [pH 7.5] for 20h. Afterwards the cells were washed six times with phosphate buffer (8 mM Na2HPO4, 1.5 mM KH2PO4, 133 mM KCl, and 0.8 mM MgCl2 [pH 7.4]). These agarose balls are melted at 70^0^C for 10 mins, diluted 15 fold with distilled H20 then digested with BsrG1 restriction enzymes. A 3ul aliquot of mock or digested sample was used for ssDNA analysis by quantitative PCR.

### H2A Ubiquitination assays

H1299 cells were co-transfected with control siRNA, HP1βsiRNA or Flag H2A plasmid. Exponentially growing cells were irradiated (10 Gy) and samples harvested at different time after irradiation. H2A ubiquitination was analyzed with anti-Flag antibody (Sigma) (Ahmed et al., 2018; Singh et al., 2018).

### Cloning and protein purification

Human FL-RING1A (1–406aa) and CSD HP1β(80–185aa) were subcloned into bacterial expression 1GFP vector (Addgene #29663) and 1B vector (Addgene #29653), respectively and proteins were purified as previously described (Hambarde et al., 2021) with changes in buffers. Briefly, RING1A was purified from HisTrap column, TEV-cleavage to remove His-GFP tags, and Superdex 75 column in the final buffer (25 mM HEPES, pH 7.6, 150 mM NaCl, 100 µM ZnCl2, 5% glycerol, 5 mM BME); CSD HP1β was purified from HisTrap column and Superdex 75 column in the final buffer (25 mM HEPES, pH 7.5, 150 mM NaCl, 2% glycerol, 1 mM DTT, 1 mM BME).

### Invitro pull down assay

FL-RING1A was incubated with His6-CSD HP1β in molar ratio 1:2 in buffer H (20 mM Tris, pH 8, 70 mM NaCl, 5% glycerol, 5 mM BME, 100 µ M ZnCl2) on ice for 20 min. The mixture was loaded onto Ni-NTA resin, which was pre-equilibrated with Buffer H and wash with buffer H excessively, followed by elution with Buffer H plus 300 mM imidazole. The fractions were analyzed by SDS-PAGE and western blot using anti-RING1A (GTX50803 from GeneTex) and anti-HP1β (sc-20699 from Santa Cruz Biotechnology) antibodies.

### Structure prediction and modelling

Full length RING1A and HP1β models were predicted by a deep learning-based method, TrRosetta (Yang et al., 2020), using webserver (https://robetta.bakerlab.org). The predicted models were evaluated by error estimate and the known crystal structure homologs [Ring1B (PDB: 4R8P, chain L); HP1β(PDB: 3F2U for CD; PDB: 5T1G for CSD)]. The best models were docked together by ClusPro protein docking webserver (https://cluspro.bu.edu) (Kozakov et al., 2017), and the top 10 ranked docking models were evaluated by known crystal structure homolog (PDB: 4R8P). RRING1B-Bmi1 interactions. Final docking model was generated by using Chimera and structural conservation was performed using ConSurf webserver (https://consurf.tau.ac.il) (Ashkenazy et al., 2016; Landau et al., 2005; Pettersen et al., 2004) by inputting the TrRosetta models of RING1A and HP1β.

## Acknowledgements

We acknowledges NIH grants supported by NIH R01 CA129537 and R01 GM109768 and The Houston Methodist Research Institute. Special thanks are due to Nobuo Horikoshi for generating cell lines. Work by JAT and CLT was supported by P01 CA092584, R35 CA220430, Cancer Prevention Research Institute of Texas (CPRIT) grant RP180813, and a Robert A. Welch Chemistry Chair. We thank Walter N Hittelman and Nitika Taneja for their suggestions.

## AUTHOR CONTRIBUTIONS

V.C., C.R.H., and T.K.P. directed the study. V.C., C.R.H., R.P., C.L.T., S.C. and T.K.P. contributed to the design. V.C., R.K.P., C.R.H., S.C. and T.K.P. performed the experiments. J.A.T. X.W. and C.L.T did in-vitro experiments, structure analysis, prediction, and modeling. V.C., C.R.H., K.T,. C.L.T., J.A.T. and T.K.P wrote the paper.

---

**Supplementary Fig. 1:**
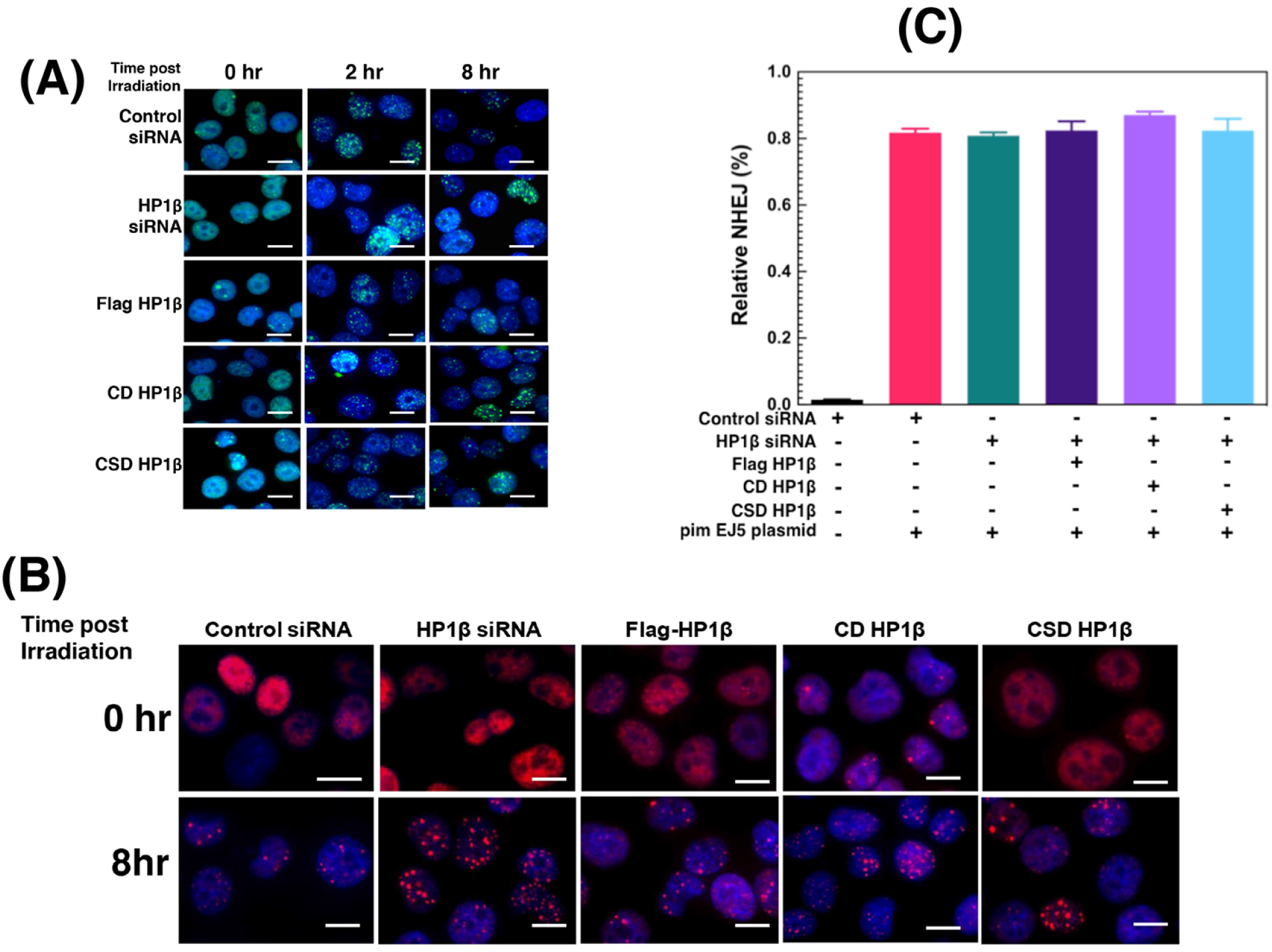
HP1β deletion mutants graphical representation and its DNA damage response. (**A&**B) Immuno fluorescent images of γ-H2AX and 53BP1 foci at different times after irradiation (2Gy). Scale bar shown here is 20μm. Cell were depleted of endogenous HP1β with siRNA then transfected with the indicted FLAG tagged HP1β expression vectors before irradiation. **C**) Non-homologous end joining DNA repair as measured by the pimEJ5 plasmid assay in HP1β depleted cells supplemented with full length or HP1β deletion mutants. 3 individual experiments were conducted and an SD value in between experiments was determined.

**Supplementary Fig. 2:**
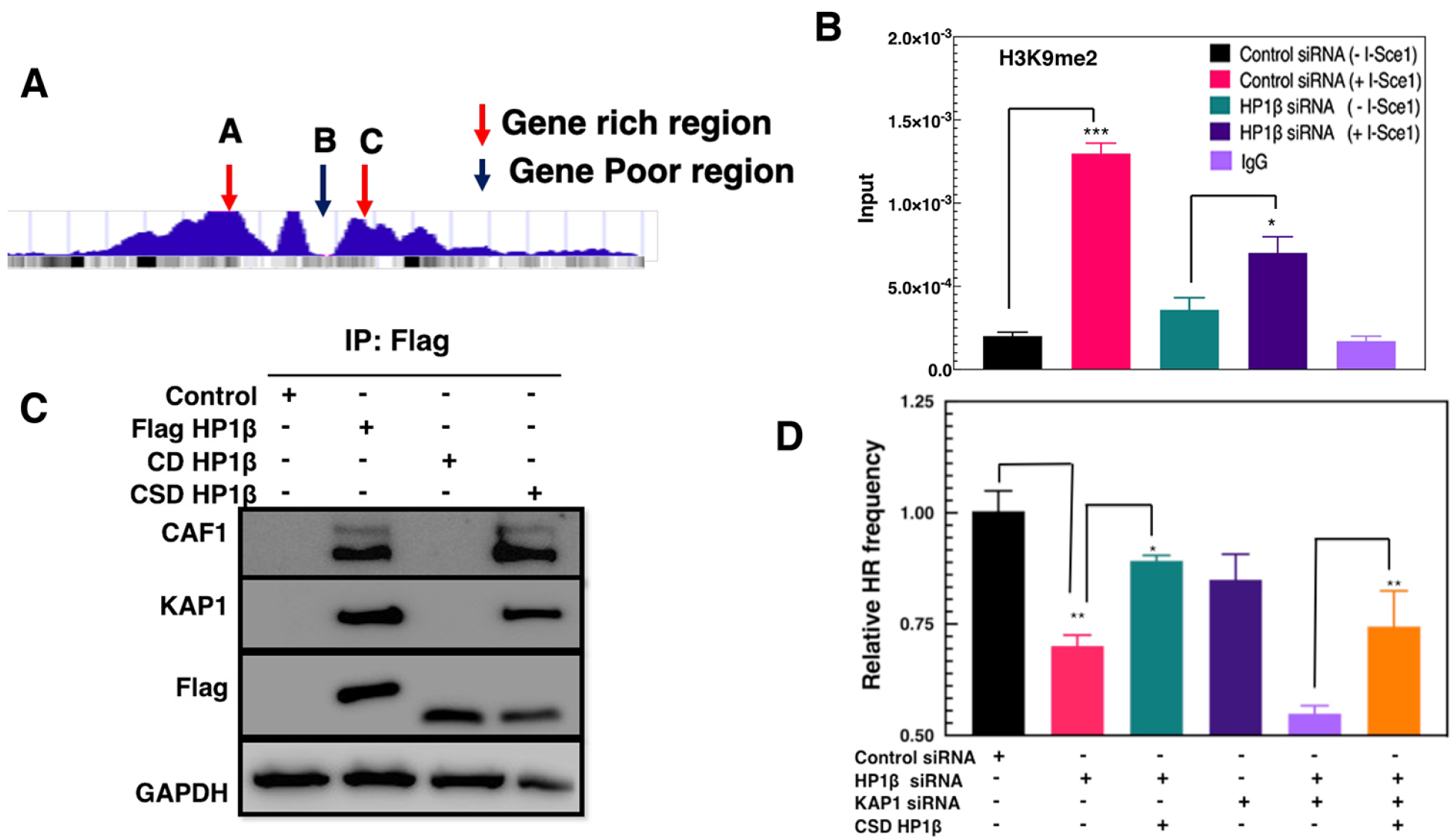
HP1β mutant recruitment to DSB sites in independent of KAP1. A) I-Sce1 containing unique sites in gene-rich and -poor regions of human chromosomes. B) Enrichment of H3K9me2 levels after I-SceI inducible DNA DSBs. C) Full length HP1β and its mutant (CDHP1β and CSDHP1β co-immunoprecipitation analysis with CAF1 and KAP1. D) Homologous Recombination in CSD HP1β expressing cells after KAP1 depletion measured by DR-GFP assay. Three to four independent experiments were carried out, SD values was determined. ^*^*p* < 0.05; ^**^*p* < 0.01; ^***^*p* < 0.001.

**Supplementary Fig. 3:**
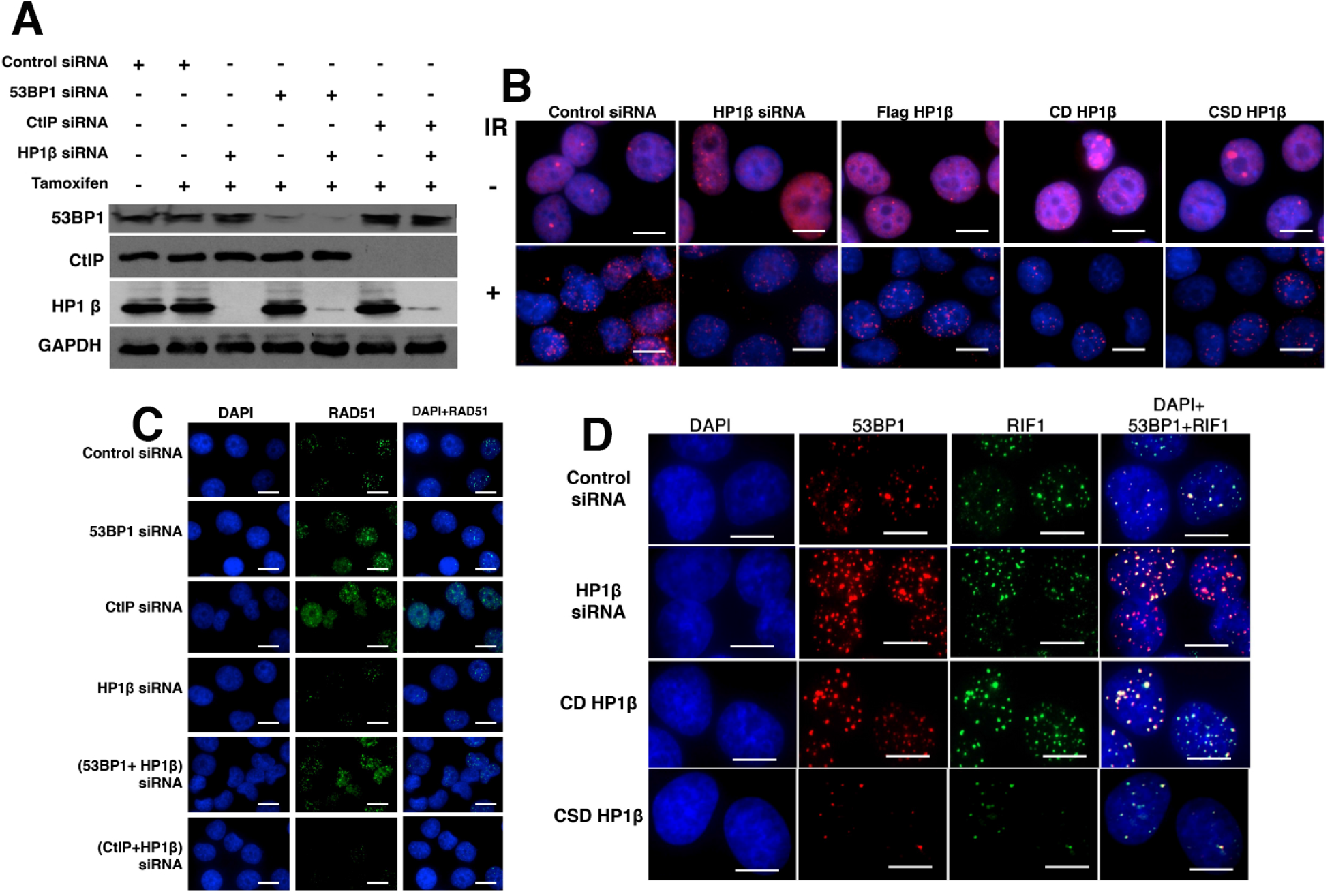
DNA end resection associated repair foci formation in irradiated HP1β depleted cells expressing deletion mutants. A) Western blot showing the depletion of HP1β, 53BP1 and CtIP with specific siRNAs. B&C) Immuno fluorescent visualization of RAD51 foci in different samples as described in Fig. 3 with and without Irradiation (10Gy). Scale bar shown here is 20 μm. D) 53BP1 and RIF1 foci formation after 4 hrs of post Irradiation in HP1β depleted cells expressing either the CD or CSD HP1β. Scale bar is 10 μm.

**Supplementary Fig. 4:**
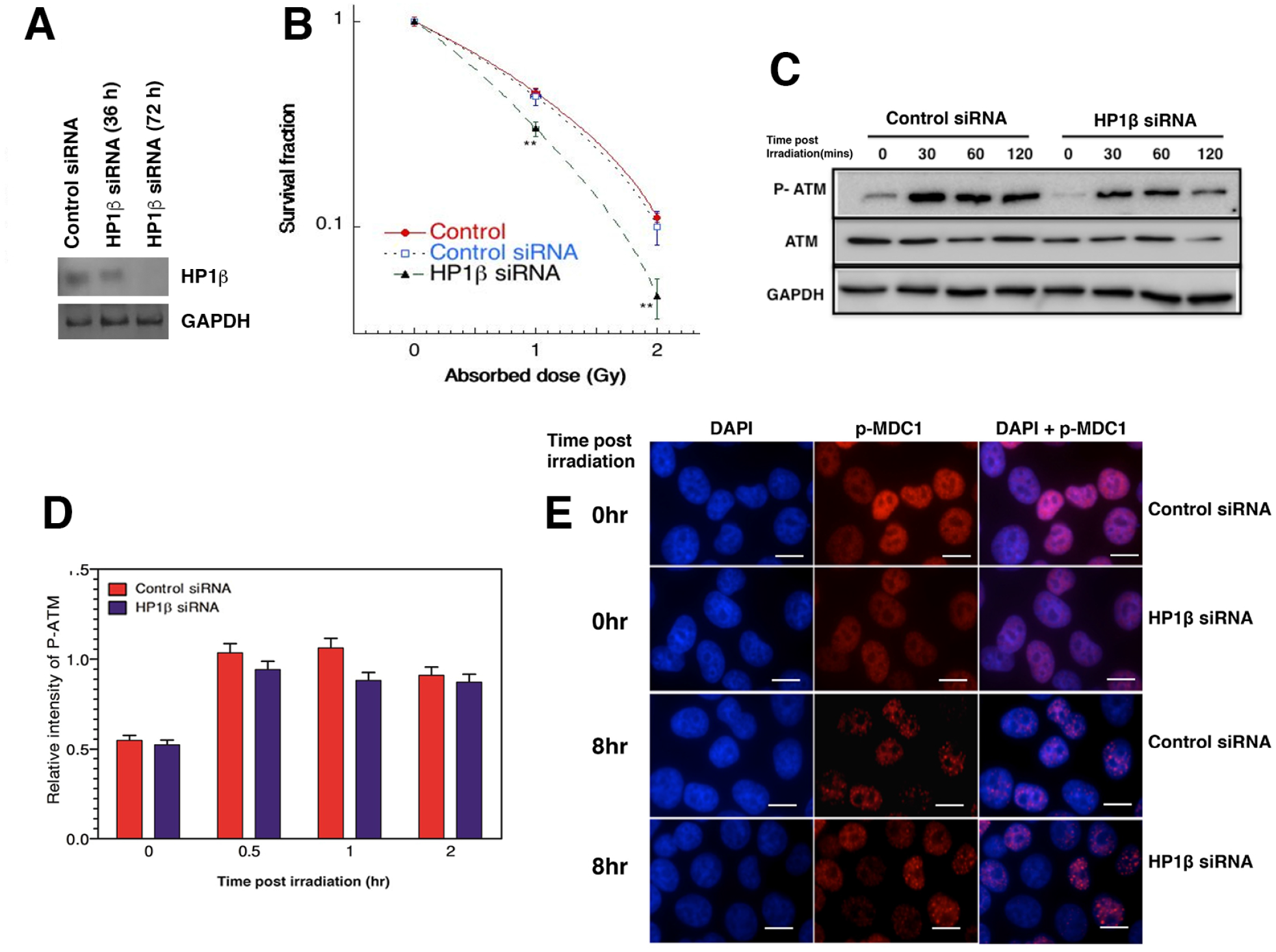
HP1β depletion has no impact on the ATM mediated DNA damage response. A) Western blot showing depletion of HP1b with specific siRNA in GM5823+hTERT cells. B) Clonogenic cell survival after IR exposure of GM5823+hTERT cells with and without HP1 depletion. B&C) Western blot analysis of cells with and without HP1β irradiated with 10Gy. Cell lysates were prepared 0, 30, 60,120 mins after radiation. D) Cells with and without HP1β were immune stained for p-MDC1 after radiation. Scale bar is 20μm. Three independent experiments were carried out and SD values determined. *p* < 0.05; ^**^*p* < 0.01.

**Supplementary Fig. 5:**
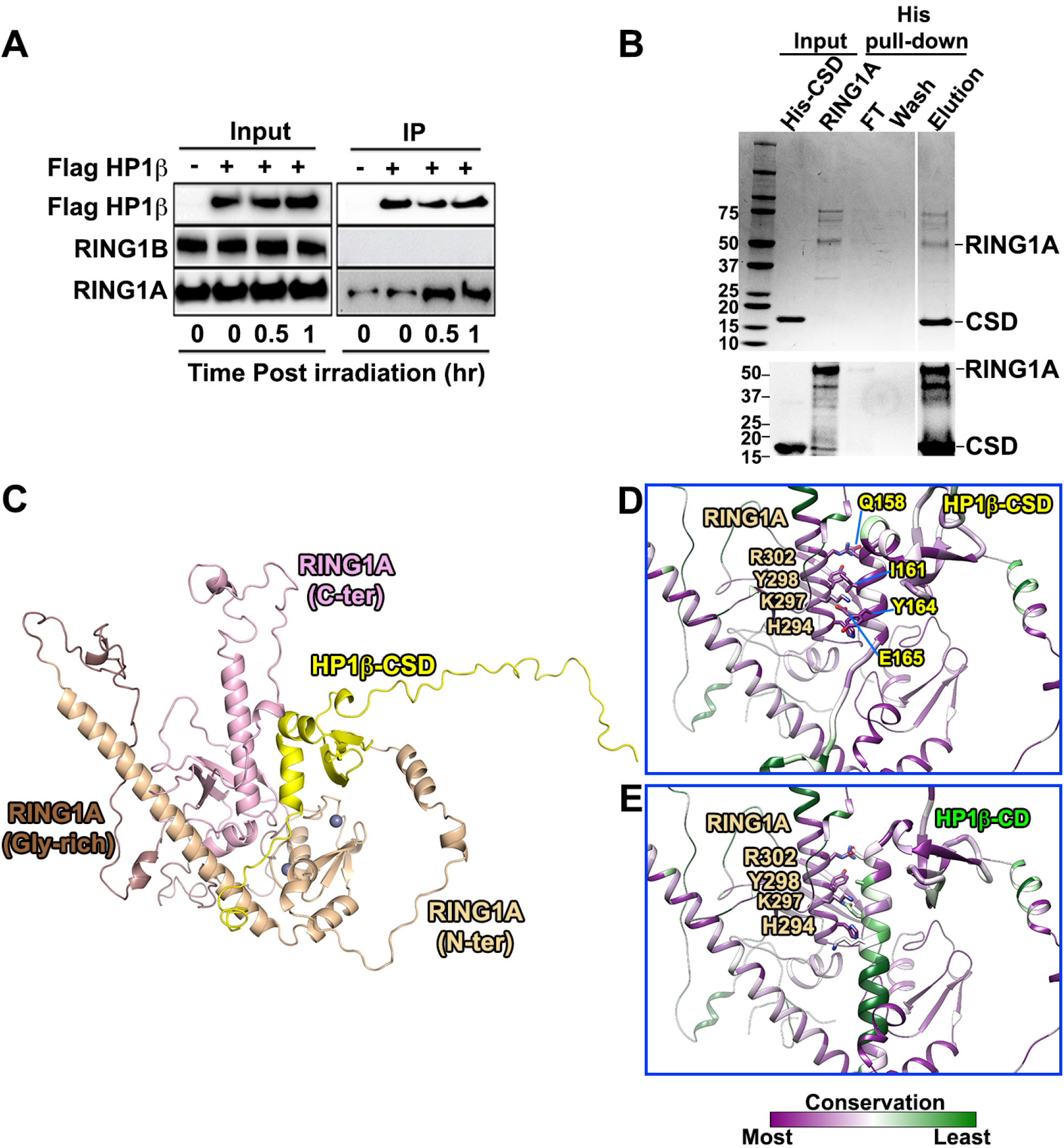
HP1β interaction with PRC1 complex protein after irradiation and Model of RING1A and CSD HP1β interaction. A) Co-immunoprecipitation analysis of endogenous RING1A and 1B with Flag HP1β, with and without irradiation). B) Invitro pull-down of recombinantly purified RING1A and CSD HP1β proteins; top, Coomassie gel; bottom, western blot. C) Rigid-body docking model of HP1β (80–185aa) with FL-RING1A. Zn ions are shown in gray spheres. D&E) Sequence conservations are mapped onto RING1A and HP1β models. The CD HP1β model is aligned over the CSD HP1β docking model. The potential interaction residues between RING1A and CSD HP1β are labelled.

